# Cholecystokinin release triggered by presynaptic NMDA receptors produces LTP and sound-sound associative memory formation

**DOI:** 10.1101/188839

**Authors:** Xiao Li, Xi Chen, Yin Ting Wong, Haitao Wang, Hemin Feng, Xuejiao Zheng, Jingyu Feng, Joewel T. Baibado, Robert Jesky, Yujie Peng, Zhedi Wang, Hui Xie, Wenjian Sun, Zicong Zhang, Xu Zhang, Ling He, Nan Zhang, Zhijian Zhang, Peng Tang, Jun-Feng Su, Ling-Li Hu, Qing Liu, Xiao-Bin He, Ailian Tan, Xia Sun, Min Li, Kelvin Wong, Xiaoyu Wang, Yi Ping Guo, Fuqiang Xu, Jufang He

## Abstract

Memory is stored in neural networks via changes in synaptic strength mediated in part by NMDA-dependent long-term potentiation (LTP). There is evidence that entorhinal cortex enables neocortical neuroplasticity through cholecystokinin (CCK)-containing neocortical projections. Here we show that a CCK_B_ antagonist blocks high-frequency stimulation (HFS)-induced LTP in the auditory cortex, whereas local infusion of CCK induces LTP. CCK^-/-^ mice lacked neocortical LTP and showed deficits in a cue-cue associative learning paradigm; administration of CCK rescued associative learning. HFS of CCK-containing entorhino-neocortical projection neurons in anesthetized mice enabled cue-cue associative learning. Furthermore, when one cue was pre-conditioned to footshock, the mouse showed a freezing response to the other cue, indicating that the mice had formed an association. HFS-induced neocortical LTP was completely blocked by either NMDA antagonist or CCK-BR antagonist, while application of either NMDA or CCK induced LTP after low-frequency stimulation (LFS). Moreover, in the presence of CCK LTP was still induced, even after blockade of NMDA receptors. Local application of NMDA induced CCK release in the neocortex. To identify how NMDA receptor switches LTP, a stimulation protocol of 25 pulse-pairs was adopted to replace HFS; NMDA-dependent LTP was induced with the inter-pulse intervals between 10 and 100 ms, but not with those of 5 and 200 ms. LTP-mediated plasticity was linked to localization of the NMDA receptor subunit NR2a on cortical CCK terminals originating in the entorhinal cortex. These novel findings suggest that presynaptic NMDA receptors on CCK terminals control the release of CCK, which enables neocortical LTP and formation of cue-cue associative memory.

**One Sentence Summary:** Presynaptic NMDA receptors switches the release of CCK from entorhinal neurons, which enables neocortical LTP and formation of sound-sound associative memory.

---

Memory is stored in neural networks through changes in synaptic strength *(1, 2)*. Long-term potentiation (LTP) and long-term depression (LTD) are two forms of synaptic plasticity that are considered to represent a neural basis of memory in different brain regions *(3-11)*. The major form of LTP in the hippocampus and neocortex is induced through tetanus burst stimulation (TBS) or high frequency stimulation (HFS) *(3-5, 10, 12)*. Previous studies have shown that NMDA receptors play a crucial role in HFS-induced LTP in the hippocampus *(13-17)* and neocortex *(18, 19)*, and in the formation and consolidation of associative memory *(20, 21)*.

Serving as the gateway from the hippocampus to the neocortex, the entorhinal cortex forms strong reciprocal connections with the neocortex *(22-24)* and shows extensive cholecystokinin (CCK) labeling *(25-27)* with projections to neocortical areas, including the auditory cortex *(22, 23, 28)*. CCK is the most abundant cortical neuropeptide *(29)*, and mice lacking the CCK gene exhibit poor performance in a passive avoidance task and display impaired spatial memory *(30)*. We previously found that local infusion of CCK into the auditory cortex of anesthetized rats induces plastic changes that enable auditory cortical neurons to start responding to a light after its pairing with an auditory stimulus *(28)*. Activation of the entorhinal cortex potentiates neuronal responses in the auditory cortex, and this effect is suppressed by infusion of a CCK_B_ antagonist *(28)*, suggesting that the entorhinal cortex enables neocortical plasticity via CCK-containing neurons projecting to neocortex.

If CCK enables cortical neuroplasticity and associative memory formation, then we would expect CCK-induced neuroplasticity to affect LTP. The release of neuropeptides occurs slowly in response to repetitive firing *(31, 32)*. Therefore, we hypothesize that HFS activates both postsynaptic neurons and presynaptic terminals, including those containing CCK. This in turn leads to CCK release and LTP induction in the neocortex. Indeed, our work has shown that CCK enables cortical neuroplasticity and associative memory formation, which correlates with the emerging insight that CCK plays a role in triggering LTP *(28)*. Thus, we further hypothesized that CCK release is controlled by presynaptic NMDA receptors.

In the present study on rats and transgenic and wildtype mice, we used optogenetic stimulation, *in vivo* extracellular recordings, *in vitro* extracellular and intracellular recording, and behavioral testing to examine: 1) the role of CCK released from the entorhinal cortex on neocortical LTP induction and cue-cue associative memory formation; 2) the relationship between CCK release and NMDA receptors; and 3) the localization of NMDA receptors on neocortical terminals of entorhinal CCK projection neurons.

## RESULTS

### Role of CCK in neocortical LTP induction

We inserted stimulation and recording electrodes in the auditory cortex of rats, targeting layers 4 and 2/3, respectively, and confirmed that TBS readily induced LTP (Fig. 1A, one-way ANOVA, F (1, 530) = 220.4, p<0.001, n=14 from 5 rats). Previously, we proposed that CCK acts as a chemical switch that enables neuroplasticity *(28)*. We hypothesized that HFS induces CCK release, which leads to cortical LTP induction. To test this hypothesis, we used ELISA to measure the CCK concentration in the cerebrospinal fluid (CSF) after TBS in the auditory cortex (left inset, Fig. 1B). CCK concentration increased from undetectable before to 5.37±1.00 pg/ml during the first 30 min after TBS in the auditory cortex (Fig. 1B, one-way ANOVA, F (2,25) = 30.2, p<0.001, n=7 from 5 rats; post-hoc before vs first 30min after the TBS p<0.001, also see Supplementary Fig.1), and recovered to 0.55±0.24 pg/ml in the second 30 min interval after TBS. To further confirm our hypothesis that CCK is the chemical switch for LTP induction, we infused a CCK_B_ receptor antagonist, L365,260, into the auditory cortex before the first TBS (Fig. 1C). We found that infusion of L365,260 blocked LTP induction, whereas infusion of vehicle had no effect on LTP induction. In L365,260-infused rats, there was no significant change in the slope of field excitatory postsynaptic potentials (fEPSPs) after the first TBS (two-way ANOVA, significant interaction, F (3,2028) =28.8, p<0.001; post-hoc p=0.182, n=10 from 8 rats). The fEPSP slope increased by 11.1% (post-hoc p<0.01, n=10) and 16.4% (post-hoc p<0.001, n=10), respectively after the second and third TBS. This modest level of LTP induction can be explained by the diffusion of L365,260 into CSF circulation more than 60 min after infusion at the second TBS, and more than 120 min after infusion after the third TBS. By contrast, in vehicle-infused rats, the fEPSP slope increased by 30.4% after the first TBS (post-hoc p<0.001, n=12 from 11 rats), 19.6% after the second TBS (post-hoc p<0.001, n=10), and 6.19% after the third TBS (post-hoc p=0.01, n=10), indicating the saturation effect for repeated TBS. After three episodes of TBS, L365,260-infused rats showed significantly less potentiation of fEPSPs than vehicle-infused rats (post-hoc *p*<0.001), suggesting that the blockade of CCK_B_ receptors on neocortical neurons suppresses LTP induction.

**Fig. 1.**
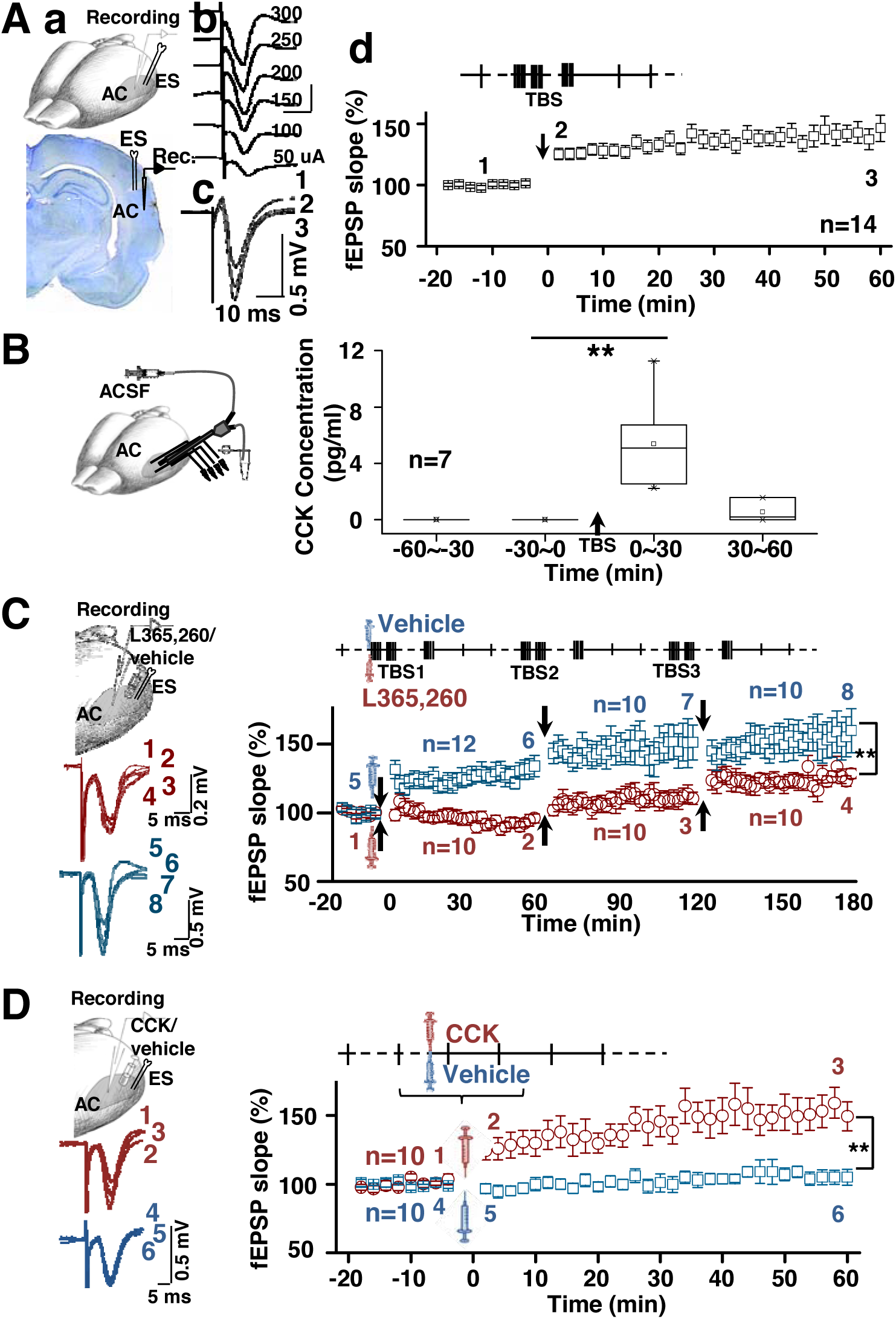
Role of CCK in neocortical LTP induction on the rats. **(A)** (a) Position of recording and stimulating electrodes in the auditory cortex of rats. (b) Representative relationship between input currents and evoked fEPSPs. (c) Representative fEPSPs before (1) and after (2, 3, corresponding to the numbers indicated in d) TBS. (d) Normalized slopes of fEPSPs before and after TBS.Stimulation paradigm above the fEPSP curve. **(B)** Left: Diagram of stimulation electrode placements and micro-dialysis. Right: A box chart showing concentrations of CCK before and after TBS in the auditory cortex based on enzyme-linked immunosorbent assay (ELISA). At the baseline level, from −60min to 0min, the concentration of CCK was lower than the detection limit (1pg/ml) of the ELISA kit, set as 0. After TBS, the concentration of CCK increased and could be measured (5.37 ± 1.15 pg/ml) (** One-way ANOVA, p<0.001), but dropped quickly in the following half an hour. **(C)** Upper left: Position of recording and stimulating electrodes and pipette in the auditory cortex of rats. Lower left: Representative fEPSPs before and after injection of L365,260 (1-4) or vehicle (5-8) and TBS. Right: Normalized slopes of fEPSPs before and after TBS with L365,260 (red circle) or vehicle (blue square) injection. TBS stimulation protocol above the curve (** Two-way ANOVA p<0.001). **(D)** Upper left: Position of recording and stimulating electrodes and pipette in the auditory cortex of rats. Lower left: Representative fEPSPs before and after injection of CCK (1-3) or vehicle (4-6). Right: Normalized slopes of fEPSPs before and after CCK (red circle) or vehicle (blue square) injection (** Two-way ANOVA p<0.001). The experimental protocol above the curve. Data are expressed as mean ± SEM.

To further test the role of CCK in cortical LTP induction, we replaced TBS with infusion of CCK in the area between the stimulating and recording electrodes (Fig. 1D). CCK infusion without TBS induced LTP in the auditory cortex of rats (two-way ANOVA showed significant interaction, F (1, 756) =86.1, p<0.001; post-hoc before vs after CCK infusion, *p*<0.001, n=10 from 6 rats), whereas no LTP was induced by vehicle infusion (two-way ANOVA, post-hoc *p*=0.43, n=10 from 2 rats).

### LTP and memory deficits in CCK^-/-^ and CCK suppressed mice

If CCK acts as a chemical switch that enables neocortical neuroplasticity, then we would expect deficits in neocortical LTP in CCK^-/-^ mice. Indeed, no LTP was induced by TBS in the auditory cortex of CCK^-/-^ mice (two-way ANOVA, significant interaction F (1,1357) = 152.1, p<0.001; post-hoc *p*=0.39, n=20 from 20 mice, Fig. 2A), whereas wildtype C57 mice showed significant TBS-induced LTP (two-way ANOVA; post-hoc *p*<0.01, n=16 from 16 mice). Importantly, CCK-/- mice neurons in the auditory cortex responded to auditory stimuli and produced event-related potentials similar to the wild type mice (Supplementary Fig. 2), indicating that the auditory pathway was functionally intact in CCK-/- mice.

**Fig. 2.**
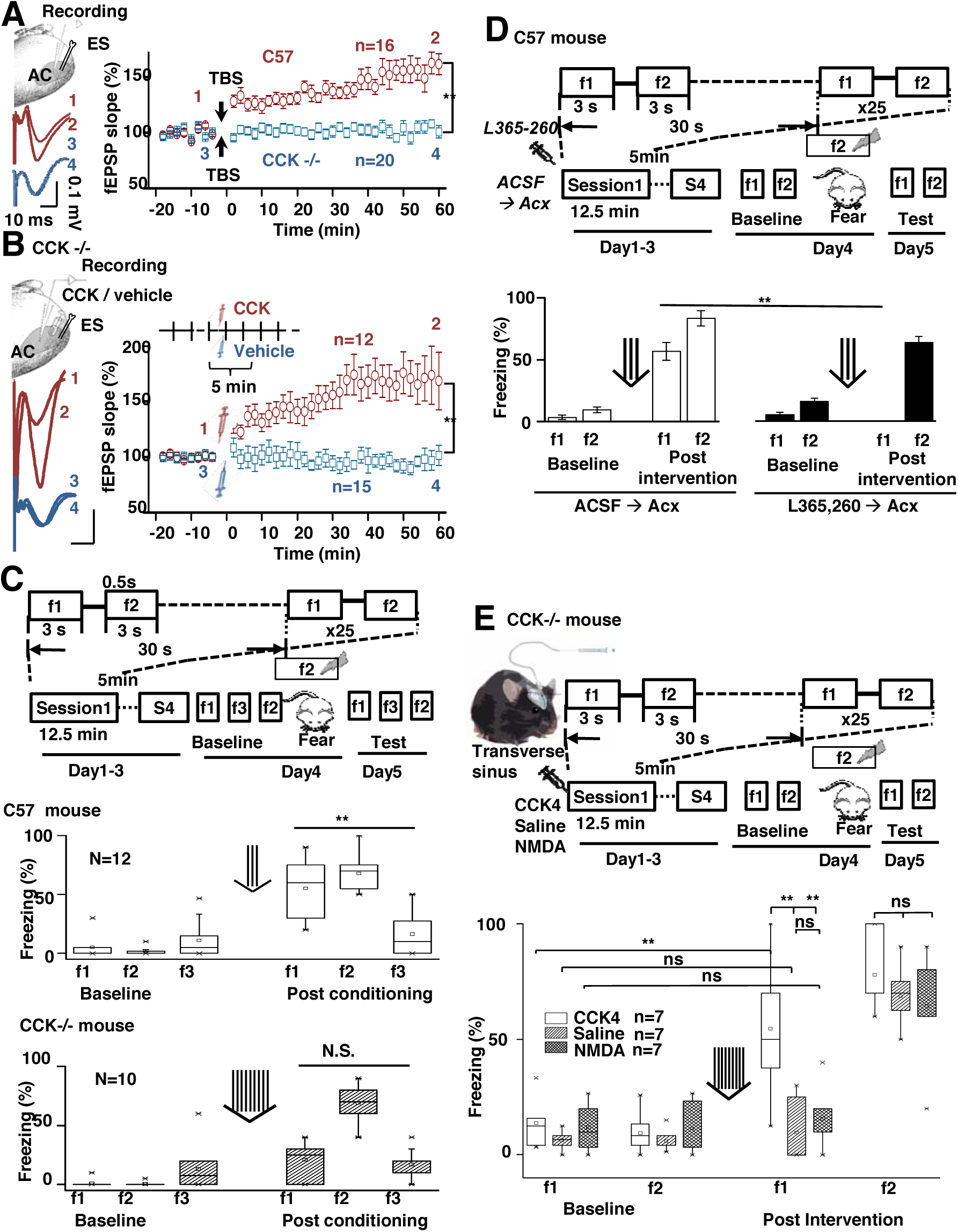
Neocortical LTP and learning an association between two tones were demolished in CCK^-/-^ mice, but rescued after administration of CCK in CCK^-/-^ mice. **(A)** Upper left: Position of recording and stimulating electrodes in the auditory cortex of mice. Lower left: Representative fEPSPs before and after TBS in C57 mice (1-2) and CCK^-/-^ mice (3-4). Right: Normalized slopes of fEPSPs before and after TBS in C57 mice (red circle) or CCK^-/-^ mice (blue square) (** Two-way ANOVA p<0.001). **(B)** Upper left: Position of recording and stimulating electrodes and pipette in the auditory cortex of mice. Lower left: Representative fEPSPs before and after injection of CCK (1-2) or vehicle (3-4). Right: Normalized slopes of fEPSPs before and after CCK (red circle) or vehicle (blue square) injection (** Two-way ANOVA p<0.001). The experimental protocol above the curve. (**C**) Upper: Diagram of a training protocol for mice to associate the tones of f1 and f2. Lower: Box charts show freezing percentages of the C57 (open) and CCK^-/-^ (shaded) to tones of f1, f2, and f3, before and after the conditioning (** Two-way ANOVA p<0.001). (**D**) Upper: Diagram of a training protocol for mice to associate the tones of f1 and f2. Lower: Box charts show freezing percentages of the C57 with ACSF (open) and C57 with L365,260 (shaded) to tones of f1and f2, before and after the conditioning (* one-way ANOVA, Tukey Test, p<0.05). (**E**) Upper: Diagram of implanted drug infusion cannula. The training protocol for mice to associate the tones of f1 and f2 was the same as above. Lower: Box charts show freezing percentages of CCK^-/-^ (shaded) mice to tones of f1 and f2, before and after the intervention (drug infusion before tones pairing; Three-way ANOVA CCK-F1 test vs baseline p<0.001, Saline-F1 test vs baseline p=0.613, NMDA-F1 test vs baseline p=0.564; test-F1 CCK vs. Saline p<0.001, CCK vs. NMDA p<0.001, NMDA vs. Saline p=0.358; there is no statistically significant difference between CCK vs Saline vs NMDA for test-F2). Data are expressed as mean ± SEM.

To determine if CCK receptors in CCK-/- mice were functional and whether infusion of CCK in the neocortex has a rescuing effect on LTP in the cortex, we infused CCK between the stimulation and recording electrodes in the auditory cortex, similar to the above mentioned rat experiments (Fig.1D). CCK infusion induced LTP in CCK-/- mice (two-way ANOVA, significant interaction F (1,984) =99.8, p<0.001; post-hoc before vs after CCK infusion *p*<0.001, n=12 from 12 mice, Fig. 2B), whereas no LTP was induced by vehicle infusion (post-hoc *p*=0.51, n=15 from 15 mice).

To determine if the lack of LTP in CCK^-/-^ mice is associated with impaired learning and memory, we tested for formation of associative memory between two auditory stimuli; i.e., cue-cue association. We assumed that mice would be able to associate the cues, after presenting the two cues (tone f1 and tone f2; 3 sec each) in sequence with an interval of 500 ms in between and a 30-sec inter-trial-interval for a total of 300 trials for three days (100 trials comprised of four 25-trial sessions per day). We then conditioned the tone f2 with footshock on day 4. We expected that the mice would show a fear response not only to tone f2, but also to tone f1, as the mouse would associate tone f1 with tone f2. In the first experiment, we compared the formation of associative memory following the paradigm noted above in CCK^-/-^ and CCK^+/+^ mice. The C57 mice showed a significantly higher freezing percentage to tone f1 (1 kHz, Movies S1,4) that was paired with f2 (4 kHz, Movies S2,5) than to tone f3 (16 kHz, Movies S3,6) that was not previously paired with f2 (55.0±8.0 % vs 16.3±5.1 %, two-way ANOVA, significant interaction F(2,66) =22.9, p<0.001; post-hoc *p*<0.001, N=12, Fig. 2C), indicating that an associative memory was formed between the f1 and f2 tones after 300 paired trials in C57 mice. While the difference between the freezing percentages to f1 and f3 was not statistically significant on CCK^-/-^ mice (21.0±5.3% vs 17.0±4.2%, two-way ANOVA, significant interaction F(2,54) =34.5, p<0.001; post-hoc *p*=0.49, N=10, Movie S7-12), the conditioning trials required for the CCK^-/-^ to achieve the same freezing percentage as the C57 mice were 3 times more (9 vs 3).

In the second experiment, we compared the influence of the infused CCK antagonist in the auditory cortex during tone f1 and f2 pairing on the formation of associative memory in C57 mice, following the same associative memory paradigm as mentioned above. Mice that received a bilateral infusion of L365,260 in the auditory cortex showed no freezing (Movies S19, 22), while those with vehicle infusion (ACSF in DMSO) showed a significant amount of freezing time when f1 was presented (56.7±7.1% vs 0±0%, one-way ANOVA, Tukey Test, p<0.05; Fig. 2D; Movies S13, 16; a third tone was adopted as another control for f1, Movies S15, 18, 21, 24; Supplementary Fig. 3). No infusion was performed during pre and post-treatment testing. As showed in Fig. 2C, CCK-/- mice displayed deficits in the formation of associative memory.

**Fig. 3.**
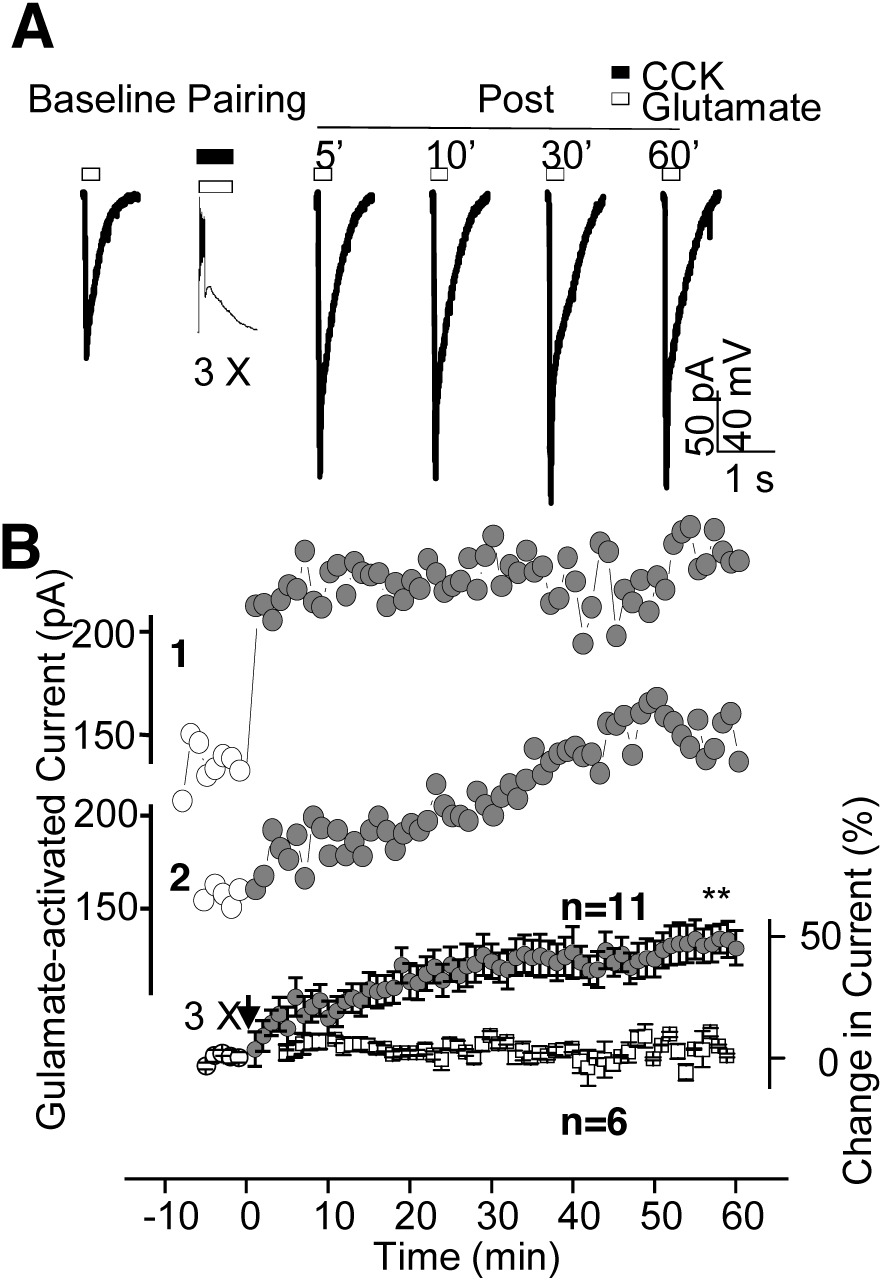
Glutamate-activated current response potentiated after a 3-trial pairing of postsynaptic depolarization and co-application of both glutamate and CCK. **(A)** Glutamate-activated current responses are shown before (Baseline) and at 5, 10, 30, and 60 min after the pairing. In this experiment, CCK was repeatedly puffed simultaneously with glutamate. The interval between two puffs was 30 s. **(B)** The time courses of the potentiation of glutamate-activated current for the above neuron and a typical neuron showing a gradually increased glutamate-activated current are shown in the upper curves before (open circle) and after (filled circle) the pairing. The time course of the change of the group data is shown in the lower curve (circle), together with the control group with no pairing (square) (** Two-way ANOVA p<0.001). Data are expressed as mean ± SEM.

In the third experiment, we infused CCK-4 through an implanted cannula in the venous sinus on the scalp of CCK-/- mice. Infusion enabled the formation of an associative memory in CCK-/- mice rescuing the associative memory deficit in CCK-/- mice. However, neither infusion of saline or NMDA (1.5mM in saline) improved the freezing time after the presentation of f1 (10.2+/-1.9% vs 46.4+/-6.3%, 6.3+/-4.0% vs 9.6+/-5.0%, 11.9+/-3.6% vs 15.7+/-4.8%, CCK4/Saline/NMDA; Three-way ANOVA CCK-f1 test vs baseline p<0.001, Movie S33,34, Saline-f1 test vs baseline p=0.613, Movie S25,26, NMDA-f1 test vs baseline p=0.564, Movie S29,30; test-f1 CCK vs. Saline p<0.001, CCK vs. NMDA p<0.001, NMDA vs. Saline p=0.358; there was no statistically significant differences of among CCK vs Saline vs NMDA treatments for f2 tests; Fig. 2E; Movie S27,28,31,32,35,36).

### Postsynaptic current potentiated after pairing neuronal firing with simultaneous puff-applications of CCK and glutamate

To determine whether LTP could be independently produced postsynaptically, glutamate puffing was adopted to directly activate the receptors at the postsynaptic membrane of cultured cortical neurons. Cultured cortical neurons showed no direct change in membrane potential to CCK puffing in a time window of tens of seconds or to repeated applications of CCK in a short time window within a second (Supplementary Fig.4). However, the neuron responded to glutamate puffing at the soma or dendrites (Baseline, Fig. 3A). Based on our previous *in vivo* intracellular recordings, three conditions including: (1) pre- and (2) postsynaptic co-activities, and (3) the presence of CCK must be fulfilled in order to produce synaptic plasticity *(28)*. Given that memories can be rapidly formed in the mammalian brain, we hypothesized that synaptic plasticity would happen after a few paring trials. To test this hypothesis, we paired depolarizing current injections of the recorded neuron with simultaneous puff-applications of glutamate and CCK for three trials (Fig. 3A). The glutamate-activated current changed from 129.9 pA before the pairing to 222.4 pA at 5 min, 223.8 pA at 10 min, 242.8 pA at 30 min, and 230.1 pA at 60 min after the pairing (Fig. 3A). The five-trial averaged responses of the neuron over the experimental course are shown in Fig. 3B (neuron 1). The majority of the neurons showed a gradual increase in the glutamate-activated current up to 30 and 50 min after the pairing, as shown in the neuron 2 (Fig. 3B). Population data showed a significant increase in glutamate-activated current after the pairing (two-way ANOVA, significant interaction F (1,943) =49.1, p<0.001; post-hoc before vs after paring *p*<0.001, n=11 neurons, Fig. 3B). For an average of 11 neurons, the glutamate-activated current increased by 39.7% at 30 min after the pairing, while simple glutamate puffing without pairing induced no potentiation in the current (two-way ANOVA; post-hoc *p*=0.69, n=6, Fig. 3B).

### HFS induces LTP in the auditory cortex to natural stimulus

HFS putatively induces release of CCK (*31*); thus, we surmised that pairing a natural auditory stimulus (presynaptic activation) with low-frequency activation of the auditory cortex (post-synaptic) after HFS of the auditory cortex would potentiate a significant neuronal response to the auditory stimulus, provided that the aforementioned prerequisite conditions were met. To test this hypothesis, HFS (5 pulses at 100 Hz) or LFS (LF; 5 pulses at 1 Hz) was followed by five pairings of an auditory stimulus (AS) and a single-pulse of electrical stimulation (ES, to produce post-synaptic activity of the auditory neurons; HF-ES/AS or LF-ES/AS) delivered at 1 Hz. This stimulation protocol was repeated four times at 10 sec intervals. Neuronal responses to the auditory stimulus were recorded for 16 min before and 60 min after the stimulation protocol. We observed greater potentiation of fEPSP slope after the HFS protocol (HF-ES/AS) than after the LFS protocol (LF-ES/AS) (two-way ANOVA, significant interaction F (1,1060) =61.4, p<0.001; post-hoc HF vs LF, *p*<0.001, n=15 from 7 rats for HF, n=13 from 6 rats for LF, Fig. 4A). The results indicate that HFS together with the pairing invoked a larger potentiation of neuronal networks enabling the induction of LTP in response to an auditory stimulus compared to LFS.

**Fig. 4.**
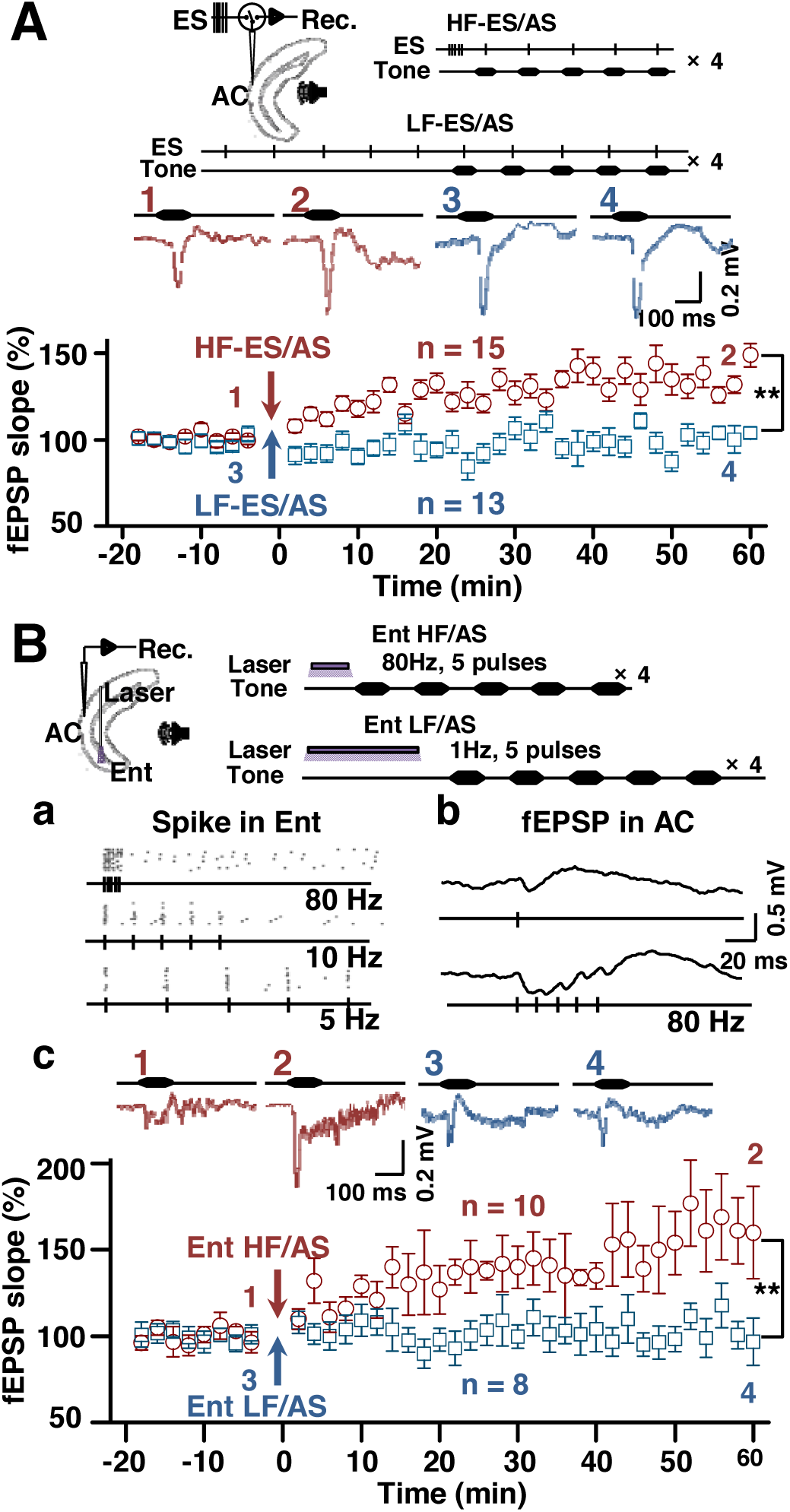
Neuronal responses to an auditory stimulus after HFS and pairings of the auditory stimulus and electrical stimulation of the auditory cortex. **(A)** Upper left: Position of shared stimulating/recording electrode in the auditory cortex of rat. Upper right: HF-ES/AS and LF-ES/AS protocols in which HFS or LFS of the auditory cortex was followed by pairings of an auditory stimulus and single-pulse electrical stimulation. Middle: Representative fEPSPs before and after HF-ES/AS (1-2) or LF-ES/AS (3-4). Lower: Normalized slopes of fEPSPs in response to the auditory stimulus before and after the HF-ES/AS (red circle) or LF-ES/AS (blue square) stimulation protocols (** Two-way ANOVA p<0.001). **(B)** Upper left: Position of the recording electrode in the auditory cortex and laser fiber in the entorhinal cortex of Thy1-Chr2-eYFP mice. Upper right: Ent HF/AS and Ent LF/AS protocols in which HFS or LFS of the entorhinal cortex was followed by presentations of an auditory stimulus. (a) Entorhinal cortex spiking responses to 80, 10, and 5 Hz laser stimulation. (b) Auditory cortex fEPSP responses to single-pulse and 80 Hz laser stimulation. (c) Upper: Representative fEPSPs before and after Ent HF/AS (1-2) or Ent LF/AS (3-4). Lower: Normalized slopes of fEPSPs in response to the auditory stimulus before and after the Ent HF/AS (red circle) or Ent LF/AS (blue square) stimulation protocols (** Two-way ANOVA p<0.001). Data are expressed as mean ± SEM.

### Entorhinal HFS induces neocortical LTP

Since entorhinal neurons that project to the neocortex enable neuroplasticity (*28*) and act as a switch for associative memory formation in the auditory cortex (*33*), we hypothesized that HFS of the entorhinal neurons would induce LTP in neural response to the auditory stimulus in the auditory cortex. We then tested this hypothesis in Thy1-ChR2-YFP transgenic mice using laser stimulation to activate entorhinal neurons (Fig. 4B). These transgenic mice express the light-activated ion channel, Channelrhodopsin-2, fused to Yellow Fluorescent Protein (YFP) under the control of the mouse thymus cell antigen 1 (*Thy1*) promoter. As all entorhino-neocortical projection neurons are CCK neurons (*28*), we expected that CCK would be released in the auditory cortex after HFS of the entorhinal neurons. We presented the noise-burst stimulus in 5 repeats at 1 Hz after HFS of the entorhinal cortex (Ent HF) in this experiment. Since the auditory neurons responded with a detectable field potential to the selected auditory stimulus, no electrical stimulation was paired with the auditory stimulus. Entorhinal neurons responded to HF laser stimulation of up to 80 Hz with spikes (Fig. 4Ba), and auditory cortical neurons responded to entorhinal HF laser stimulation with excitatory field potentials (Fig. 4Bb). HF (5 pulses at 80 Hz) or LF (5 pulses at 1 Hz; Ent LF) laser stimulation of the entorhinal cortex was followed by five presentations of an auditory stimulus at 1 Hz; this stimulation protocol was repeated four times at 10 sec intervals. Neuronal responses to the auditory stimulus were recorded for 16 min before and 60 min after the stimulation protocol. We observed that the HFS protocol (Ent HF/AS) (two-way ANOVA, significant interaction F (1,675) =29.4, p<0.001; post-hoc before vs after HF, p<0.001, n=10 recordings from 9 mice), but not the LFS protocol (Ent LF/AS) induced LTP to the auditory stimulus (post-hoc p=0.67, n=8 recordings from 7 mice, Fig. 4Bc). The results of these two experiments confirmed that projections from the entorhinal cortex are important for neocortical LTP induction and that burst-firing of entorhinal neurons may be the key to triggering CCK release and inducing LTP in the auditory cortex.

### Entorhinal projections enable neocortical LTP and associative memory

To obtain direct evidence that HFS of the entorhino-neocortical projections causes CCK release in the auditory cortex and that the released CCK enables the potentiation of neuronal responses and associative memory formation in the auditory cortex, we conducted two parallel experiments that allowed for direct optogenetic manipulation of specific projection neurons by: (1) injecting AAV-DIO-ChR2-eYFP into the entorhinal cortex of CCK-ires-Cre mice and adopting CCK^-/-^ mice (CCK-CreER) as the control group (Fig. 5) and (2) injecting AAV-CaMKIIa-ChR2(E123T/T159C)-mCherry into the entorhinal cortex of C57 mice (Supplementary Fig.5). The AAV-DIO-ChR2-eYFP is a CRE-dependent virus that only transfects CCK neurons in the injected area. The CCK-ires-Cre knock-in allele has an internal ribosome entry site and Cre recombinase in the 3' UTR of the CCK locus. As such, Cre recombinase expression is directed to CCK-expressing cells by the endogenous promoter elements of the CCK locus.

**Fig. 5.**
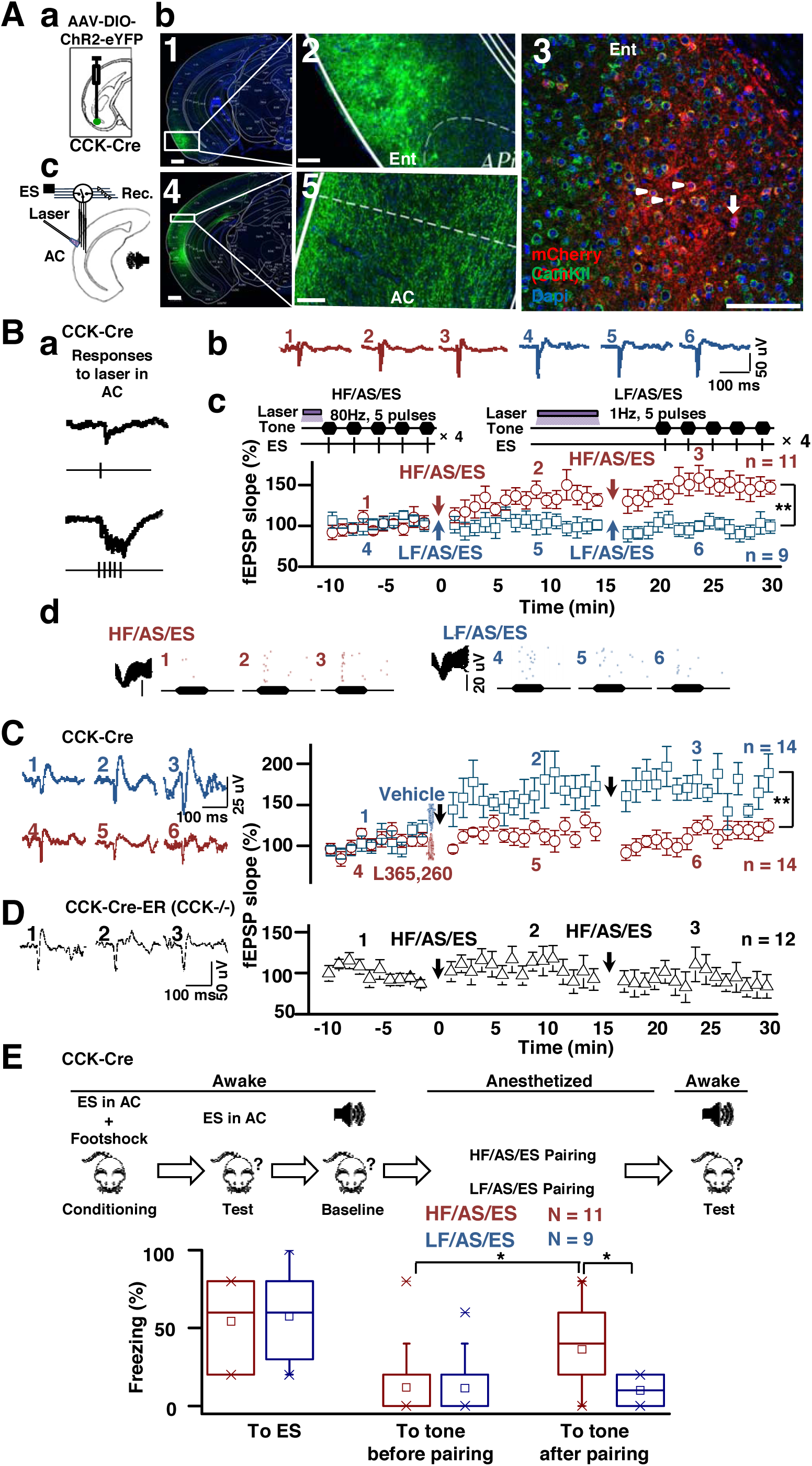
HFS of CCK-containing entorhino-neocortical projections enables the association between an auditory stimulus and electrical stimulation of the auditory cortex, leading to behavioral changes. **(A)** (a) AAV-DIO-ChR2-eYFP was injected into the entorhinal cortex of CCK-Cre mice. (b) Images of virus expression in the entorhinal cortex (Ent, 1-3) and auditory cortex (AC, 4-5) (scale bars: 500 for 1 and 4; 100 μm for 2, 3, and 5). In 3, mCherry (CCK), CamKII, and DAPI were overlapped (Arrow head: neurons express both CamkII and CCK; arrow: a neuron only expresses CCK). (c) Position of the laser fiber and stimulating/recording electrodes in the auditory cortex. **(B)** (a) Responses in field potential to laser stimulation in the auditory cortex, (b) Representative fEPSPs before and after the HF/AS/ES (1-3) and LF/AS/ES (4-6) protocols. (c) Normalized slopes of fEPSPs after the HF/AS/ES (red circle) or LF/AS/ES (blue square) stimulation protocols (** Two-way ANOVA p<0.001). (d) Unit responses to the auditory stimulus before and after the pairings of HF/AS/ES (1-3) and LF/AS/ES (4-6). (**C**) HF-stimulation induced LTP in the auditory response blocked by CCK antagonist. Left: Representative fEPSPs before and after vehicle (1-3) or CCK antagonist (4-6) injection. Right: Normalized slopes of fEPSPs after the HF/AS/ES stimulation protocols with CCK antagonist (red circle) or vehicle (blue square) injection (** Two-way ANOVA p<0.001). **(D)** No potentiation was induced in CCK^-/-^ mice with the same protocol as in **b-c**. Left: Representative fEPSPs before and after the HF/AS/ES protocol. Right: Normalized slopes of fEPSPs after the HF/AS/ES protocol. **(E)** Upper: Cued fear conditioning and stimulation protocols. Lower: A box chart shows time spent freezing in response to the paired auditory stimulus before and after the HF/AS/ES (Red) or LF/AS/ES (Blue) pairing (* Two-way ANOVA p<0.01). Data are expressed as mean ± SEM.

In the first experiment, we tested whether cortical LTP induction via HFS of entorhino-neocortical projections occurs via CCK release by injecting AAV-Dio-ChR2-eYFP into the entorhinal cortex of CCK-ires-CRE mice. Entorhinal neurons transfected with the virus projected to neocortical areas, including the auditory cortex (Fig. 5Ab). We found that most of the transfected CCK neurons in the entorhinal cortex were glutamatergic neurons (76.6% of CCK neurons (red) expressed CamKII (green), Fig. 5Ab-c). At least 6 weeks after AAV-DIO-ChR2-eYFP injection into the entorhinal cortex (Fig. 5Aa), recording/stimulating electrodes and optical fibers were implanted in the auditory cortex of CCK-ires-CRE mice (Fig. 5Ac). We observed that neurons in the auditory cortex responded to single pulses or a burst (80 Hz) of laser stimulation with excitatory field potentials (Fig. 5Ba). Next, we used a similar pairing protocol as in the previous experiment (Fig. 4B), but with additional electrical stimulation presented at 50 ms after the onset of each tone stimulus. We observed that neurons in the auditory cortex responded to single pulses or a burst (80 Hz) of laser stimulation (Fig. 5Ba). The reintroduced single-pulse electrical stimulation of the auditory cortex was used as a cue in the following behavioral test. The auditory cortical neurons showed potentiated responses to the auditory stimulus after the HF/AS/ES protocol (two-way ANOVA, significant interaction F (2,713) =29.4, p<0.001; post-hoc before vs after 1^st^ HF/AS/ES, p<0.001; after 1^st^ vs 2^nd^ HF/AS/ES, p<0.001, n=11 from 11 mice), but not after the LF/AS/ES protocol (post-hoc before vs after 1^st^ LF/AS/ES, p=0.65; after 1^st^ vs 2^nd^ LF/AS/ES, p=0.54, n=9 from 9 mice; Fig. 5Bb-c). Despite relatively short monitoring of the changes in neuronal responses (15 min after the first pairing and another 15 min after the second pairing), the pattern of potentiation was resembled that observed after the entorhinal HF/AS protocol in Fig. 4Bc. Likewise, changes in unit firing rates as indicated in the raster plots showed that the HF/AS/ES protocol lead to an increase in spike numbers, while the LF/AS/ES condition did not. Spike numbers increased in points 2 and 3 compared to 1 for the HF/AS/ES condition, while it did not increase in 5 and 6 compared to 4 for the LF/AS/ES condition (Fig. 5Bd).

Our subsequent experiments aimed to confirm whether LTP induced by HFS of the CCK entorhino-neocortical projections in the auditory cortex was generated by release of CCK. We found that vehicle infusion (ACSF) before HF/AS/ES induced LTP, while infusion of a CCK_B_ antagonist (L365,260) before HF/AS/ES failed to induce LTP in the auditory cortex of CCK-ires-Cre mice (Fig. 5C, two-way ANOVA, significant interaction F (2,1044) =16.4, p<0.001; vehicle, post-hoc before vs after 1st HF/AS/ES p<0.001, before 1^st^ vs after 2^nd^ HF/AS/ES p<0.001, n=14 from 4 mice; L365,260, post-hoc before vs after 1^st^ HF/AS/ES, p=0.205, before 1^st^ vs after 2^nd^ HF/AS/ES p=0.396, n=14 from 4 mice).

An additional control experiment was carried out on CCK^-/-^ mice, adopting the same protocol that was used in the CCK-ires-Cre mice. In CCK^-/-^ mice, CCK-CreER knock-in allele both abolishes CCK gene function and expresses the CreERT2 fusion protein in both interneurons and pyramidal neurons as directed by the endogenous CCK promoter/enhancer elements. Results showed that no potentiation was induced after the HF/AS/ES protocol (one-way ANOVA, F(2,488)=4.6, p=0.011; post-hoc before vs after 1^st^ HF/AS/ES, p=0.308; before 1st vs after 2^nd^ HF/AS/ES p=0.088, n=12 from 7 mice; Fig. 5D), suggesting that the loss of LTP induction in CCK^-^ /- mice was generated by the absence of CCK release.

We next examined whether the LTP induced by HF/AS/ES pairing in the anesthetized mouse could be demonstrated in a behavioral context. We paired electrical stimulation of the auditory cortex with footshock for 5 days until the mice showed stable freezing responses to the conditioned single-pulse electrical stimulation in the absence of footshock (Fig. 5E, To ES). An auditory stimulus that triggered no freezing response and evoked weak neuronal responses was then selected as the pairing auditory stimulus for each mouse. Pairings of either HF/AS/ES or LF/AS/ES was conducted while mice were in anesthetized conditions. The neurons that showed potentiated responses to the auditory stimulus were presumably a subpopulation of those activated by electrical stimulation. Therefore, we expected that activation of these neurons would induce freezing behavior, as electrical stimulation was previously conditioned to a footshock. Indeed, mice in the HF/AS/ES group showed more freezing in response to the auditory stimulus after pairing than before pairing (two-way ANOVA, significant interaction F (2,89) =3.2, p=0.047; post-hoc before vs after HF/AS/ES pairing p=0.002, N=11 mice, Fig. 5E), whereas mice in the LF/AS/ES group showed no change in freezing after pairing (post-hoc p=0.84, N=9 mice, Fig. 5E). Mice in the HF/AS/ES group also showed more freezing than mice in the LF/AS/ES group after pairing (36.4±6.21% vs 10.0±7.28%, post-hoc p=0.007, Fig. 5E).

To further verify these findings, we conducted a parallel experiment that included injecting AAV-CaMKIIa-ChR2(E123T/T159C)-mCherry into the entorhinal cortex of C57 mice. Results emulated those above, wherein the HF/AS/ES group showed more freezing than the LF/AS/ES group (Supplementary Fig.5). Both experiments demonstrate that an association between an auditory stimulus and electrical stimulation of the auditory cortex, reflected by potentiated neuronal responses to the auditory stimulus, could be established, while mice were under anesthesia. Moreover, pre-conditioning the electrical stimulation to a footshock caused mice to exhibit freezing in response to the auditory stimulus, providing further evidence of an association between auditory and electrical stimuli. Therefore, HFS of CCK-containing entorhino-neocortical projections appears to be a key contributor to neocortical neuroplasticity and associative memory formation.

### Either CCK or NMDA antagonist blocks HFS-induced LTP

HFS induced neocortical LTP is largely NMDA-receptor dependent. To establish a possible connection between NMDA and CCK associated LTP, we employed *in vitro* electrophysiological recordings in cortical slices from C57 mice. We began by using HFS in conjunction with common antagonists for both NMDA and CCK_B_ receptors (i.e., DL-2-amino-5-phosphonovaleric acid (APV) and L365,260, respectively). We found that HFS induced LTP in cortical slices of the C57 mouse was completely blocked by application of either NMDA or CCKBR antagonist (APV or L365,260; Fig. 6A).

**Fig. 6.**
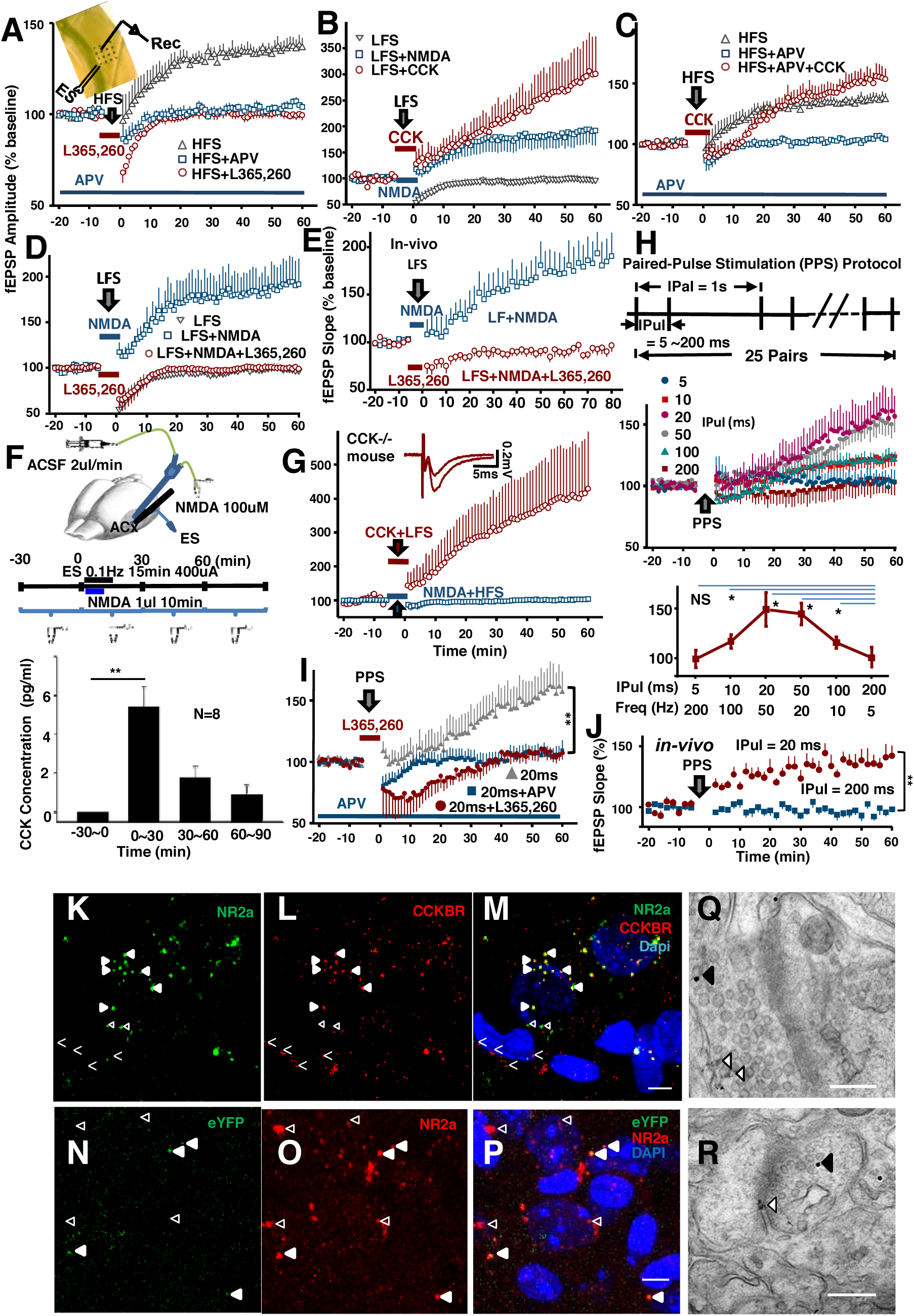
Presynaptic NMDA receptors control the release of CCK, which in turn enables CCK-dependent neocortical LTP. **(A)** Either an NMDA antagonist or CCK antagonist blocked HFS-induced LTP in C57 cortical slices. APV was applied throughout all the recording period, but L365,260, was applied after the baseline recording, and 4 min before the HFS. **(B)** Either NMDA or CCK induced LTP in C57 cortical slices with LFS. **(C)** Blocking of NMDA receptors did not block LTP in the presence of CCK in C57 cortical slices. **(D)** NMDA induced LTP was blocked by application of CCK antagonist L365,260 in the C57 slices. **(E)** NMDA induced LTP was blocked by application of CCK antagonist, L365,260, in the auditory cortex of anesthetized C57 mouse. L365,260 was dissolved in 1% DMSO + 99% ACSF. L365,260 or control (DMSO+ACSF) was infused during the whole experiment. **(F)** Local application of NMDA together with HFS triggered CCK release in the auditory cortex of C57 mouse. Micro-dialysis and ELISA was same as that in Fig. 1. **(G)** No LTP was induced by NMDA application, but a marked induction of LTP was generated by CCK application in cortical slices of CCK-/- mouse. **(H)** A new paradigm for LTP induction: 25 paired-pulses. Upper: 25 pairs of paired-pulses with varied inter-pulse interval (IPuI) between 5 ms and 200 ms and fixed inter-pair interval (IPaI) of 1 sec. Middle: Field EPSP showed the induction of LTP with different IPuIs. Lower: Potentiation of the fEPSP amplitude as a function of the IPuI. The IPuI was converted to frequency, though there were only two pulses. One-way ANOVA was applied to compare mean change of fEPSP amplitude (% baseline) of the last 10 min of IPuI = 200ms with other data points (*P<0.001). **(I)** LTP induced with the PPS protocol was blocked by infusion of either APV or L365,260. **(J)** LTP was induced by the PPS protocol when IPuI was 20 ms, but not 200 ms in the auditory cortex of anesthetized C57 mouse. **(K-M**) Co-labeling of NR2a subunits (green) **(K**), CCKBR (red) **(L**), and DAPI (blue) **(M**). Filled arrowheads indicate the co-localization of the NR2a subunits and CCKBR. Open triangles indicate labeling of NR2a subunits without co-localized CCKBR, while open arrowheads indicate the labeling of CCKBR without co-localized NR2a subunits. The images were overlapped by 12 confocal microscopic pictures of 1-μm steps; individual images are shown in SFig. 8. Scale bar: 10 μm. **(N-P**) Co-labeling of eYFP (green) **(N**), NR2a (red) **(O**) and DAPI (blue) **(P**) in the auditory cortex of CCK-Cre mouse with AAV-Dio-eYFP-ChR2-injected in the entorhinal cortex. Filled arrowheads SS indicate the colocalization of the NR2a and eYFP. Open triangles indicate the labeling of NR2a without co-localized eYFP. The images were overlapped by 12 confocal microscopic pictures with 1-μm steps; individual images shown in SFig. 9. Scale bar: 10 μm. **(Q-R**) Immuno-electron microscopy showed the co-localization of NR2a subunits in the cortical projection terminals of the entorhinal CCK neurons. Gold-particles of 15 nm show the immunoreactivities to mCherry, while those of 6 nm show the immunoreactivities to NR2a subunits. AAV-Dio-mCherry was injected in the entorhinal cortex of CCK-Cre mouse. Closed arrowheads indicate the 15-nm particles, while open triangles indicate the 6-nm particles. Scale bar: 200 nm.

### LTP can be induced with LFS in the presence of CCK or NMDA

Normally, LFS does not induce LTP in cortical slices as represented in fEPSP. Our initial recording reaffirmed this. However, infusion of either NMDA or CCK in conjunction with LFS for 5 min produced significant LTP (two-way ANOVA, F(2,1494) = 57.73, P<0.001;increased by −0.84%+/-4.56%, n=6, post-hoc before vs after the LFS P>0.05; increased by 90.29%+/-25.01%, n=7, post-hoc before vs after the LFS with NMDA p<0.001;, increased by189.35%+/-67.85%, n=7, post-hoc before vs after the LFS with CCK p<0.001 Fig. 6B). To further evaluate how NMDA and CCK may interact to produce LTP, we explored whether a CCK_B_ antagonist could occlude NMDA-dependent LTP induction.

### NMDA-dependent LTP can be blocked by CCK antagonist

Based on the amplitude difference in the LTP induction with infusion of NMDA and CCK respectively (Fig. 6B), it was predicted that CCK was the direct switch for LTP induction, while NMDA was assumed to control the release of CCK. To verify this hypothesis, LTP induction was examined with HFS and with both HFS and CCK after blocking NMDA receptors with APV. We found that induction of LTP was generated upon application of CCK even when NMDA receptors had been blocked, supporting our hypothesis (Fig. 6C). If NMDA is critical to the release of CCK, a CCK_B_ receptor antagonist would interrupt the binding of CCK at the postsynaptic receptor site, resulting in lack of blockade of NMDA-associative LTP. Consistent with this hypothesis, co-application of NMDA and L365,260 completely blocked LFS/NMDA induced LTP (Fig. 6D). When the same experimental conditions were repeated in the auditory cortex of anesthetized mice, we found that NMDA-induced LTP was blocked when co-applied with L365,260 (Fig. 6E).

### No rescuing effect of NMDA application on LTP and cue-cue association of CCK-/- mice

The earlier *in vivo* experiment showed that local application of CCK induced LTP in the auditory cortex of CCK-/- mouse. Similarly, we found that in cortical slices of CCK-/- mice application of CCK lead to LTP (Fig. 6G). However, NMDA application with pairing of HFS in cortical slices of CCK-/- mice failed to produce LTP (Fig. 6G). In earlier behavioral experiment on CCK-/- mice, the tone-tone association was not improved after intravenous infusion of NMDA (Fig. 2E). These results indicate that NMDA receptors are not acting after CCK receptor in LTP induction, and likely on CCK release. Taken together, this implies that LTP induction stems from NMDA-mediated release of CCK. To advance this hypothesis investigated the mechanisms of CCK release.

### CCK release is controlled by NMDA receptors

We used micro-dialysis and ELISA to establish that local application of NMDA induced CCK release in the neocortex (Fig. 6F). The CCK concentration in the dialyzed CSF increased significantly from 0 (undetectable level) to 5.38±1.04 pg/ml (one-way ANOVA, F (3, 28) = 13.18, p<0.001, n=8; post-hoc before vs first 30min after TBS, p<0.001), and returned to 1.75±0.24 pg/ml in the following 30 min and 0.89±0.49 pg/ml in between 60 and 90 min.

### Paired-pulse stimulation (PPS) protocol instead of HFS protocol to induce NMDA-dependent LTP

The difference between the high-and low-frequency stimulation stems from the interval between two consecutive pulses: one is short and the other one is long. We hypothesized that the first stimulation activates the glutamatergic CCK terminal to release glutamate that then activates the presynaptic NMDA receptors; whereby the activated NMDA receptors will induce CCK release, as long as the second stimulation arrives within the critical interval. To verify this hypothesis, a new stimulation protocol was employed to replace the HFS protocol for LTP induction: 25 pulse-pairs (inter-pair-interval, IPaI = 1s; inter-pulse-interval within the pulse-pair, IPuI, varied from 5 to 200 ms; see the upper panel for the paradigm, Fig. 6H). The PPS protocol of IPuI between 10 and 100 ms induced LTP, while that of IPuI of 5 ms and 200 ms did not (Fig. 6H). A confirmation of the PPS proposal for LTP was performed with the *in vivo* preparation, which showed that PPS of IPuI of 20 ms induced LTP in the auditory cortex in anesthetized mice, while the IPuI of 200 ms did not (30.5±1.0 % vs −2.4±1.0 %, F (1, 794) = 119.2, P<0.001, two-way ANOVA, Fig. 6J). These results show that a paired-pulse stimulation of 50 to 20 Hz generated an optimal time interval between 20 and 50 ms, whereby the potentiations of fEPSP amplitudes were 56.73%+/-18.27% and 51.81%+/-11.63%, respectively (two-way ANOVA, F(1,1121) = 0.576, P>0.05;n=8, post-hoc before vs after the PPS of IPuI in 20ms p<0.001; n=7, post-hoc before vs after the paired-pulse stimulation of IPuI in 50ms p<0.001, Fig. 6H). Next, in efforts to reveal how the required initial stimulation of presynaptic NMDA receptors may affect CCK release, we applied APV during the paired-pulse stimulation. This completely blocked the potentiation (two-way ANOVA, F(1,121) = 33.738, P<0.001; potentiations of fEPSP amplitudes were 0.12%+/-1.59%n=7, post-hoc before vs after the PPS of IPuI in 20ms with APV perfusion p>0.05 Fig. 6I). Similarly, application of CCKBR antagonist (L365,260) completely blocked the potentiation (two-way ANOVA, F(1,1121) = 48.29, P<0.001 potentiations of fEPSP amplitudes were −7.39%+/-1.70% n=7, post-hoc before vs after the PPS of IPuI in 20ms with L365,260 perfusion p>0.05 Fig. 6I).

In other words, the first action potential generates glutamate release, which in turn activates NMDA receptors creating the conditions necessary for CCK release. The results imply that the critical interval for the second action potential to release CCK from the terminal falls between 10 and 100 ms. This mechanism also explains why LTP could be induced with LFS, when NMDA was applied: with presynaptic NMDA receptors activated, each pulse of LFS is able to release CCK that induces LTP (Fig. 6D). Co-application of CCKBR antagonist prevents this LTP induction (Fig. 6D). The conjoint electrophysiological data strongly suggested a concerted interaction between NMDA and CCK. Therefore, we set out to clarify whether or not NMDA receptor subunits are co-expressed with CCK.

### Localization of NR2a subunits in entorhinal CCK terminals in the auditory cortex

To evaluate relevant anatomical positioning of NMDA receptors and CCK, we performed immunohistochemical experiments and found through both confocal and electron microscopy that: 1) NR2a-containing NMDA receptors are closely co-localized with CCKB receptors (Fig. 6K-L, Supplementary Fig. 8); and 2), the projections from the entorhinal cortical CCK neurons to the auditory cortex of AAV-Dio-eYFP labled CCK-Cre mice were co-localized with NR2a subunits as evidenced by confocal (Fig. 6N-P, Supplementary Fig. 9) and electron microscopy (Fig. 6Q and R).

## DISCUSSION

The present study demonstrates that CCK plays an important role in neocortical LTP induction and associative memory formation. Local infusion of a CCK_B_ antagonist into the auditory cortex of anesthetized rats blocked HF stimulation-induced LTP, whereas infusion of CCK into the cortex was sufficient to induce LTP without HF stimulation. CCK^-/-^ mice exhibited a lack of HF stimulation-induced LTP as well as deficits in the associative learning paradigm of two tones. Cortical application of the CCK antagonist L365,260 blocked the formation of associative memory in C57 mice, while intravenous infusion of CCK-4 enabled the formation of associative memory in CCK-/- mice. The glutamate-activated current was significantly potentiated after pairing the depolarizing current injection into the recorded neuron with simultaneous puff-applications of glutamate and CCK on cultured cortical neurons. Although there are different sources of CCK that project to the auditory cortex, we focused on projections from the entorhinal cortex. We hypothesized that HFS of CCK-containing entorhinal projection neurons can potentiate neuronal responses in the auditory cortex. To test this hypothesis, we applied HFS of the auditory cortex before pairing auditory and electrical stimuli, which induced LTP to the auditory stimulus. Next, we found that HFS of the entorhinal cortex or entorhino-neocortical projections also potentiated neuronal responses to the paired auditory stimulus. Pre-application of a CCK antagonist blocked the HFS-induced potentiation. We confirmed that HFS of CCK-containing entorhinal projection neurons enabled neocortical neuroplasticity by pre-conditioning the electrical stimulation to footshock, which caused mice to freeze in response to the paired auditory stimulus.

We explored the relationship between NMDA-dependent LTP and CCK dependent LTP. In cortical slices, either a CCK or NMDA antagonist blocked HFS-induced LTP. In the presence of CCK or NMDA, LTP was induced with LFS. LTP was induced with CCK application even when NMDA receptors were blocked. Both *in vivo* and *in vitro* cortical slice experiments showed that co-application of a NMDA and CCK antagonist completely blocked the LFS/NMDA induced LTP. The micro-dialysis and ELISA experiments additionally confirmed that local application of NMDA induced CCK release in the neocortex. In CCK-/- mice, it was found that there was no induction of LTP after NMDA application nor when it was coupled with HFS. Likewise, in CCK-/- mice, attempts to rescue the formation of tone-tone associative memory after intravenous infusion of NMDA could not be obtained.

Finally, we examined the link between NMDA activation and TBS/HFS in induction of LTP with a new protocol of 25 paired-pulses. We adopted a new stimulation protocol of PPS, and found that LTP was induced with 25 pairs of IPuI between 10 and 100 ms and that was blocked by the application of APV. Anatomically, we found that eYFP labeled terminals in the auditory cortex of CCK-Cre mouse are co-localized with NMDA receptor subunit NR2a, and that NR2a subunits are generally co-localized with CCK_B_ receptors.

LTP, which was discovered in the hippocampus over 40 years ago (*3, 4*), has generally been regarded as the synaptic basis of learning and memory in vertebrates (*7*). Indeed, a recent study that selectively manipulated synapses with HFS and LFS successfully demonstrated that LTP was linked with associative memory (*11*). Our results complement those of recent studies using optogenetics to examine how neuronal assemblies support memories (*11, 34*-*36*). We found that LTP in the rat and mouse auditory cortex was induced by HFS of the auditory cortex, CCK infusion, or HFS of entorhino-neocortical projections. Together, our results lead us to conclude that HFS likely causes entorhinal projection neurons to release CCK into the auditory cortex, thereby inducing LTP in the auditory cortex in response to pairing an auditory stimulus with electrical stimulation.

In additional experiments, we dissected the different components of neocortical LTP induction. Using patch-clamp recordings of cultured cortical neurons, we concluded that in the presence of CCK, LTP can be produced postsynaptically with just a few pairings of the pre- and post-synaptic activities. Furthermore, LTP was induced by CCK infusion into the auditory cortex without HF stimulation, while no LTP could be induced in CCK^-/-^ mice or in the presence of a CCK antagonist. Based on the hypothesis that HFS induces CCK release in the cortex, we tested whether the potentiation of neuronal responses occurs only for HF-stimulated inputs. After HFS of the auditory cortex, we observed LTP in response to a probe auditory stimulus with low frequency presentation. These results imply that LTP also occurs for non-HF-stimulated inputs, extending our current understanding that LTP only occurs for HF-stimulated inputs *(37-39)*. Furthermore, HFS of CCK-containing entorhino-neocortical terminals before the pairing of auditory and low frequency electrical stimuli led to LTP to the auditory stimulus, indicating the importance of CCK in inducing neocortical LTP. Also, when the electrical stimulation was conditioned to a footshock before the pairing, the auditory stimulus triggered freezing responses in mice, indicating that mice had formed an association between electrical and auditory stimuli that meaningfully affected behavior.

Consistent with previous findings that CCK is associated with learning and memory *(30, 40-44)*, we found that CCK^-/-^ mice showed deficits in associative memory formation and cued fear conditioning tests. Although most studies have focused on GABAergic CCK neurons *(45)*, previous works indicated that many CCK neurons in the neocortex are excitatory (46, 47). In the present study, CCK-containing entorhinal neurons in CCK-Cre mice labeled with AAV-DIO-eYFP-ChR2 projected all the way to the auditory cortex and other neocortical areas. Laser stimulation of these terminals elicited fEPSPs in the auditory cortex, suggesting that they are glutamatergic neurons. Furthermore, our results suggest that memory enhancement by deep-brain stimulation of the entorhinal cortex *(48)* may be related to the activation of CCK-containing entorhinal neurons. Formation of paired visual associations or paired visuoauditory associations critically depends on the perirhinal and entorhinal cortex *(33, 49)*, yet CCK^-/-^ mice showed deficits in associating two different tones. The AS/ES pairing after HF activation of the entorhino-neocortical CCK terminals established the artificial association between the auditory and electrical stimuli in the present study. Association of the artificial manipulation using electrical stimulation and natural stimulus suggests there lies within neuroengineering and therapeutic applications.

Previous studies have shown that NMDA receptors enable the formation of associative memory and precipitate LTP induction (*13*-*18, 21*). Interestingly, cortical application of a CCK antagonist blocked the formation of the association of two tones with different frequencies in the C57 mice. Moreover, CCK^-/-^ mice exhibited deficits in the formation of the above association, and while infusing NMDA failed to rescue the formation of associative memory, intravenous infusion of CCK-4 proved effective in alleviating these deficits. These results thus imply that CCK acts as a switch for the formation of associative memory, and is seemingly more vital in this scenario than NMDA receptors. Much of the earlier aspects of this study focused on postsynaptic NMDA receptors. In our cortical slice experiments, the required conditions that generated LTP suggested that the activation of presynaptic NMDA receptors of glutamatergic CCK terminals controls the release of CCK, which enables plastic alterations in the postsynaptic neuron. Our brain slice experiments also unveiled that the TBS/HFS in LTP induction can be replaced with paired-pulses with short intervals between the pulses within each pair. Concomitantly, the data suggests that the first pulse within the pair activates the glutamatergic CCK terminals to release glutamate that in turn activates presynaptic NMDA receptors, which then act as a timer requisite for switching CCK release. Importantly, data suggest that CCK is released only if the second pulse arrives within the critical interval (10 to 100 ms).

Finally, our immunohistochemical experiments with fluorescence imaging and EM imaging confirmed that the neocortical projecting terminals of entorhinal CCK neurons in the auditory cortex are co-localized with NMDA receptor subunits (i.e., NR2a). This anatomical foundation aids the substantiation of our hypothesis that presynaptic NMDA receptors are primarily responsible for triggering the release of CCK in the neocortex. Importantly, this is congruent with previous findings that demonstrated that NR2a subunits are located in presynaptic terminals (*50*).

In summary, evidence suggests that the presynaptic NMDA receptors of CCK terminals control the release of CCK. This enables CCK-dependent neocortical LTP and the formation of cue-cue associative memory formation. To the best of our knowledge this is the first set of results showing a potentially new mechanism that underpins associative memory formation through neuropeptidergic modulation and entorhinal stimulation.

## ACKNOWLEDGEMENTS

We thank Guoping Feng and Minmin Luo for sharing of CCK transgenic mouse lines for our preliminary study, Eduardo Lau for administrative and technical assistance, Tomas Hökfelt (Karolinska Institutet), Richard Salvi (University at Buffalo), Kuanhong Wang (NIH), Robert Oswald (Cornell), and Bin Hu (Calgary) for critical comments, and Micky Tortorella (Guangzhou) Colin Blakemore (University of London), and Longnian Lin (Shanghai) for insightful discussion. This work was supported by Hong Kong Research Grants Council, National Key Basic Research Program of China, Natural Science Foundation of China, and Health and Medical Research Fund (2012CB966300, 2013CB530900, 561212M, 561313M, 11101215M, 11166316M, C1014-15G, T13-607/12R, 31200852, 03141196, 01121906, 31171060, 31371114, 31571096). We also thank the following charitable foundations for their generous support: Charlie Lee Charitable Foundation, Fong Shu Fook Tong Foundation, and Croucher Foundation.

## AUTHOR CONTRIBUTION

JFH, XL, XC, FQX designed the experiments; XL, XC, WJS, JYF, JFS, YTW, XYW collected and analyzed the data of in-vivo LTP; XC, XJZ, JYF, LH, NZ, ZJZ, WJS, ZCZ, PT, JFS, QL, XBH, YJP, ALT, collected and analyzed the data of optogenetics including physiology and behavioral experiments; YTW, HX, XZ, collected and analyzed the data from brain slices; HTW, LLH collected and analyzed the data from cultured neurons; HMF, XJZ, YJP, XL, ZDW, KW, ML collected the data of behavioral experiments; JYF collected the micro-dialysis data; RJ, XJZ, JTB, XS, XC collected and analyzed the anatomy data; YPG and FQX prepared some experimental setup; JFH, XL, and XC wrote the manuscript.

## Supplementary Materials

Materials and Methods

Figures S1-S9

Movies S1-S36

References (*51-55*)

